# Heat Shock Protein 27 versus Estrogen Therapy for Post-Menopausal Atherosclerosis: Rethinking Mechanisms of Cholesterol Lowering

**DOI:** 10.1101/2020.06.04.135483

**Authors:** Nadia Maarouf, Yong-Xiang Chen, Chunhua Shi, Jingti Deng, Catherine Diao, Matthew Rosin, Vipul Shrivastava, Zarah Batulan, Jingwen Liu, Edward R. O’Brien

**Affiliations:** Department of Cardiac Sciences, Libin Cardiovascular Institute of Alberta, University of Calgary Cumming School of Medicine, Calgary, Alberta, Canada; Department of Veterans Affairs Palo Alto Health Care System, Palo Alto, California

**Keywords:** Atherosclerosis, Cholesterol, PCSK9, LDL Receptor, Estrogen, Menopause

## Abstract

**Aims:** The estrogen-inducible protein Heat Shock Protein 27 (HSP27) as well as anti-HSP27 antibodies are elevated in healthy subjects compared to cardiovascular disease patients. Vaccination of *ApoE*^*-/-*^ mice with recombinant HSP25 (rHSP25, the murine ortholog), boosts anti-HSP25 levels and attenuates atherogenesis. As estrogens promote HSP27 synthesis, cellular release and blood levels, we hypothesize that menopause will result in loss of HSP27 atheroprotection. Hence, we now compare the efficacy of rHSP25 vaccination *vs*. estradiol (E2) therapy for the prevention of post-menopausal atherogenesis.

**Methods and Results:** *ApoE*^*-/-*^ mice subjected to ovariectomy (OVX) showed a 65% increase atherosclerotic burden compared to sham mice after 5 weeks of a high fat diet. Relative to vaccination with rC1, a truncated HSP27 control peptide, atherogenesis was reduced by 5-weekly rHSP25 vaccinations (−43%), a subcutaneous E2 slow release pellet (−52%) or a combination thereof (−82%). Plasma cholesterol levels declined in parallel with the reductions in atherogenesis, but relative to rC1/OVX mice plasma PCSK9 levels were 52% higher in E2/OVX and 41% lower in rHSP25/OVX mice (p<0.0001 for both). Hepatic LDLR mRNA levels did not change with E2 treatment but increased markedly with rHSP25 vaccination. Conversely, hepatic PCSK9 mRNA increased 148% with E2 treatment *vs*. rC1/OVX but did not change with rHSP25 vaccination. In human HepG2 hepatocytes E2 increased PCSK9 promoter activity 303%, while the combination of [rHSP27 + PAb] decreased PCSK9 promoter activity by 64%.

**Conclusion:** The reduction in post-OVX atherogenesis and cholesterol levels with rHSP25 vaccination is associated with increased LDLR but not PCSK9 expression. Surprisingly, E2 therapy attenuates atherogenesis and cholesterol levels post-OVX without altering LDLR but increases PCSK9 expression and promoter activity. This is the first documentation of increased PCSK9 expression with E2 therapy and raises questions about balancing physiological estrogenic / PCSK9 homeostasis and targeting PCSK9 in women – are there effects beyond cholesterol?

## Introduction

Our laboratory is interested in understanding why atherosclerosis is relatively silent in women compared to men – at least until menopause (MP) when estrogen levels plummet. Noting the relative abundant presence of estrogen receptors in the artery wall, we embarked on an objective discovery project looking for estrogen receptor associated proteins that might help explain important biological differences. These studies led to the discovery that Heat Shock Protein 27 (HSP27) is as an estrogen receptor beta associated protein. The relationship between HSP27 and estrogen is complex, as HSP27 acts as a repressor of estrogen-mediated transcription *in vitro*, yet its expression and extracellular release are also partially regulated by estrogens (e.g., there is an estrogen response element found in the HSP27 promoter) ^1, 2^. Importantly, our group ^1^ and four others (using objective proteomic approaches) ^3-6^ showed that HSP27 expression is inversely corelated with the degree of coronary artery plaque burden.

Accordingly, we demonstrated in a series of experiments that augmenting HSP27 levels reduces atherogenesis ^7-10^. Moreover, when *ApoE*^*-/-*^ mice that over-express HSP27 are subjected to ovariectomy (OVX) HSP27 levels are lower and athero-protection is lost, unless estrogenic therapy is administered ^2^. Clinically, elevated HSP27 blood levels are associated with a lower 5-year risk of myocardial infarction, stroke or cardiovascular death ^8^. In women who undergo premature surgical MP because of an inherited risk of cancer, we note that HSP27 levels drop in the first post-operative year ^11^.

Interestingly, natural IgG auto-antibodies to HSP27 (AAbs) are detectable in human blood, and recently we demonstrated how HSP27 immune complexes (ICs) form, dock at the cell membrane engaging with Toll-like receptor 4 (TLR4) and compete with LPS to reduce inflammatory signaling ^12^. Via NF-κB pathway activation HSP27 ICs increase the expression of the low density lipoprotein receptor (LDLR) ^13^. Unlike the effect of statins, the effect of HSP27 on LDLR expression is divergent from the sterol regulatory element binding protein 2-mediated dual regulation of LDLR and Proprotein Convertase Subtilisin/Kexin type 9 (PCKS9), the negative regulator of cholesterol metabolism. Furthermore, vaccination of mice with rHSP25 (murine ortholog of human HSP27), augments anti-HSP25 antibodies, reduces atherogenesis by markedly decreasing total plasma cholesterol levels and increases LDLR expression ^13^. Interestingly, with rHSP25 vaccination PCSK9 levels fall, and while this may in part be due to reduced expression, it is more likely due to increased PCSK9 clearance by abundant LDLRs ^13^.

Therefore, in this study we sought to determine if rHSP25 vaccination can combat atherogenesis post-OVX, employing a surgical model of MP in *ApoE*^*-/-*^ mice and using estradiol (E2) as a comparator. *In vitro*, we also explore the effects of the HSP27 IC *vs*. E2 on the expression of PCSK9. Our results indicate that rHSP25 vaccination and E2 therapy are both effective in attenuating the early stages of atherogenesis. However, we were surprised to find that E2 increases PCSK9 transcription without altering that of LDLR, while HSP27 therapy lowers cholesterol (and PCSK9) by increasing LDLR expression.

## Materials and Methods

### Recombinant protein preparation

Plasmids encoding HIS-tagged full-length HSP27 were used to express the following recombinant proteins: i) the mouse ortholog rHSP25, or the C-terminal, biologically inactive (rC1) fragment of HSP27 (AA93-205) (**Fig. 1A**). For *in vitro* studies involving human hepatocytes, we also generated rHSP27, as previously described ^14^. Using a pET-21a vector, with the plasmids were transformed into an *Escherichia coli* expression strain (DE3). Recombinant proteins were purified with a Ni-NTA resin and Q-Sepharose™ (GE Healthcare). Endotoxin was removed by Pierce High-Capacity Endotoxin Removal Resin (ThermoFisher Scientific, Waltham, MA, USA). The purity of the final recombinant proteins was determined to be more than 99% by sodium dodecyl sulfate and polyacrylamide gel (SDS-PAGE) with an endotoxin concentration lower than 2 units/mg protein measured by Limulus Amebocyte Lysate PYROGENT™ 125 Plus (Lonza). In previous experiments we demonstrated that these recombinant proteins were functionally devoid of significant endotoxin contamination, as the addition of polymyxin B had no effect on the ability of these proteins to activate NF-κB ^9, 15^.

**Fig. 1.**
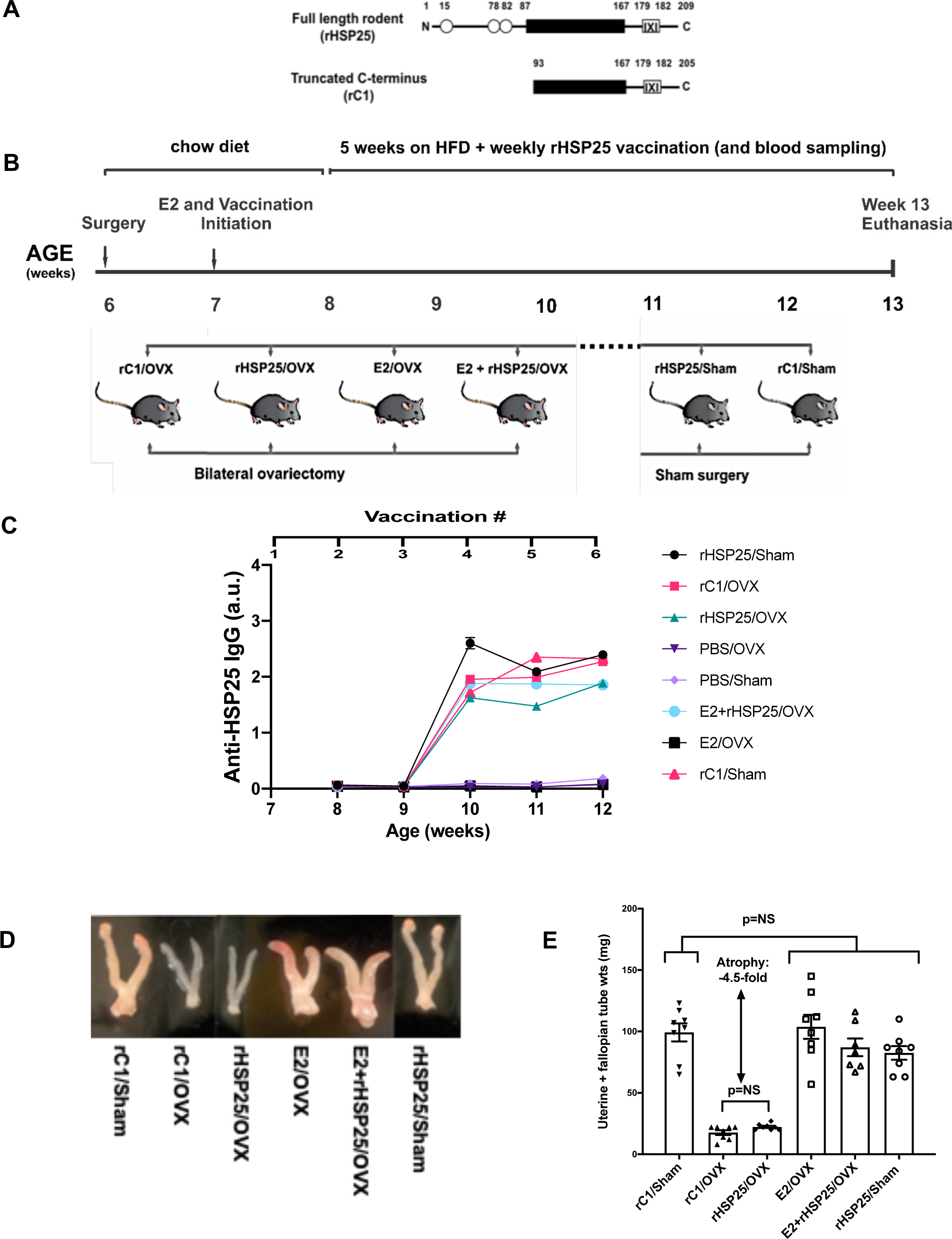
rHSP25 vaccination to prevent atherogenesis in ovariectomized *ApoE*^*-/-*^ mice. A) Schematic representation of rHSP25 (mouse ortholog of HSP27) and the biologically inactive truncated C-terminal of HSP27 (rC1 control) proteins. The black box denotes the alpha beta-crystallin domain that is important for protein oligomerization. The IXI box at the C-terminus is a flexible domain involved in the formation of multiple inter-subunit interactions. B) Age matched, sexually mature female *ApoE*^*-/-*^ mice were subjected to either OVX or sham operation (no OVX). One week later, they were randomized to the following treatments: rC1/OVX, rHSP25/OVX, E2/OVX, rHSP25+E2/OVX, rHSP25/sham and rC1/sham. A HFD was initiated one week later and continued for 5 weeks until euthanasia. There were 7-8 mice per group. See Supplemental Fig. 1 and 2 for additional control mice data. C) Anti-HSP25 IgG levels increased in all mice vaccinated with either rHSP25 or rC1. The initiation of rHSP25 vaccination began at 7 weeks of age (one-week post-surgery) and measurements were taken every week thereafter, rising to a plateau by 3-4 weeks after the first vaccination and continuing at this level until euthanasia. Data are displayed as a.u. for OD @ 450 nm. D) Representative photographs of uteri and fallopian tubes from each treatment groups. E) Combined uterine and fallopian tube weights at time of euthanasia. After OVX there was marked atrophy of the uteri and fallopian tubes of mice in the rC1/OVX and rHSP25/OVX groups compared to rC1/sham mice. There was no difference in weights between the rC1/sham and the other groups (E2/OVX, [E2 + rHSP25/OVX], rHSP25/sham), thereby suggesting that the E2 supplementation was in a physiological hormonal range. One-way ANOVA with Tukey’s multiple comparison’s test.

### Murine Surgical Model of Menopause and Atherogenesis

The following animal protocol was approved by the Animal Care and Use Committee at the University of Calgary and is in compliance with the Canadian Council on Animal Care and the NIH Guide for the Care and Use of Laboratory animals (**Fig. 1B**). Five-week-old *ApoE*^*-/-*^ female mice were purchased from the Jackson Laboratories (JAX®, Canada; C57BL/6 genetic background, Jackson Laboratory, Bar Harbor, ME, USA; stock 002052: B6.129P2-*ApoE*^*tm1Unc*^/J) and after one week of acclimatization were randomized to bilateral OVX *vs*. a surgical sham operation (i.e., ovaries not removed). For all surgical procedures, mice were given a mixture of isoflurane/oxygen (2.5%, 1L/min respectively). Briefly, the dorsum of each mouse was incised and both ovaries were removed. Muscle and skin layers were closed with sterile synthetic bio-absorbable sutures (Ethicon, Somerville, NJ). Buprenorphine (0.05 mg/kg subcutaneously) was administered twice-daily for 3 days post-surgery for analgesia. For the mice randomized to receive E2 supplementation, after recovering for one-week post-OVX a pellet containing 0.1 mg of E2 (60-day release formulation; Innovative Research America) was implanted subcutaneously between the ear and shoulder using a trocar. This strength of E2 pellet was selected because in a pilot experiment using higher concentration E2 pellets (0.25 mg/60-day) there was a 3-fold increase in the uterine / fallopian tube weight in the E2/OVX *vs*. rC1/OVX mice. Mice were also randomized to rHSP25 or rC1 vaccination, as well as the combination of rHSP25 vaccination plus an E2 pellet. Vaccination was performed weekly with recombinant HSP25 (rHSP25; 100 µg, 75 µL) or ‘rC1’, a truncated HSP27 peptide consisting of the C-terminal half of the peptide, (100 µg, 75 µL) mixed with an aluminum hydroxide gel adjuvant (Alhydrogel® 2%, Al, 25 µL; Invivogen vac-alu-250, San Diego, CA). One week after the commencement of E2 and/or vaccinations, the mice began a 5-week atherogenic High Fat Diet (HFD) containing: 1.25% cholesterol and 15.8% fat, (ENVIGO Harlan Teklad diets) until euthanasia. In addition, control experiments were conducted comparing PBS *vs*. rC1 treatments in sham and OVX mice (**Supplemental Fig. 1A**).

Mice were housed (n=3 or 4 per cage) in a temperature controlled (20°C) facility with a 12 h light/ dark cycles and had free access to autoclaved food and water. A total of 64 mice were randomized to 8 groups (8 mice/group) and all but one mouse completed the study (due to a surgical mishap). Plasma samples were collected weekly and animal weights recorded prior to treatment administration. At euthanasia, mice were placed under anesthesia with isoflurane/oxygen mixture (5%, 1L/min) and adequate sedation was verified by toe and/or tail pinching to ensure absence of a withdrawal reflex response before the mice were exsanguinated. The thoracic cavity was opened by cutting the ribs lateral to the sternum to expose the heart, and whole blood was collected by inserting a 25-gauge needle into the left ventricle. After blood collection, the right atrium pierced to permit hemorrhage, and systemic perfusion fixation with phosphate-buffered saline (PBS) thrice or until the liver turned pale in color, followed by 10% NBF. The heart and aorta were dissected, removed and immersed in NBF. The uteri and fallopian tubes were also harvested, dissected of all fat and connective tissue, photographed and weighed in order to assess the end organ effect of E2 *vs*. rHSP25 therapies.

### Murine HSP25 and anti-HSP25 IgG antibody levels

Blood samples were collected weekly prior to treatment and levels of IgG AAb were measured using an ELISA developed in our laboratory (**Fig. 1C**). Briefly, NUNC maxisorp plates (ThermoFisher) were coated with rHSP25 at a concentration of 500 ng/well in carbonate-bicarbonate buffer overnight (ON) at room temperature. The wells were blocked with 1% BSA/PBST, washed in phosphate buffered saline tween (PBST), and incubated with plasma at a final dilution of 1:2,000 in 1% BSA for two hrs followed by 3 more washes in PBST. A horse radish peroxidase (HRP) labeled goat anti-mouse IgG (H&L) antibody (catalogue #115-035-062; Jackson Immunoresearch, West Grove, PA) was used as a detection antibody at a dilution of 1:5,000 and incubated for one hr at room temperature. Finally, substrate solution (3,3’,3.5’-Tetramethylbenzidine Liquid Substrate, TMB; Millipore Sigma; Oakville, ON) was added to each well and incubated for 10 mins avoiding direct light. The reaction was stopped by 2N H_2_SO4 and the optical density quantified at 450 nm, using Synergy Mx plate reader (BioTek). To establish an internal standard for the measurement of HSP25 auto-antibodies, plasma from a healthy control subject was diluted 500 times in 1% BSA/PBST and arbitrarily defined as 50 absorbance units (a.u.).

### Evaluation of *en face* Atherosclerotic Lesion Area

The aorta was dissected from the ascending to the thoracic segments ending at the 4^th^ bronchial branches and the adventitia was removed. The primary incision followed the lesser curvature of the arch opening up the aorta longitudinally. A second incision was made along the greater curvature of the arch down to the level of the left subclavian artery. The aortic lipid content was evaluated using oil red O staining (described previously) ^10^ and photographed using an Olympus SZ-61 stereo microscope with an Olympus UC30 camera (aortic magnification ×12; **Fig. 2A, Supplemental Fig. 1B**). The *en face* atherosclerotic aortic lesions were analyzed using Image-Pro software (Media Cybernetics) to calculate the total and atherosclerotic lesion areas. The extent of atherosclerosis was expressed as the percentage of surface area of the entire aorta covered by atherosclerotic lesions ^8^.

**Fig. 2.**
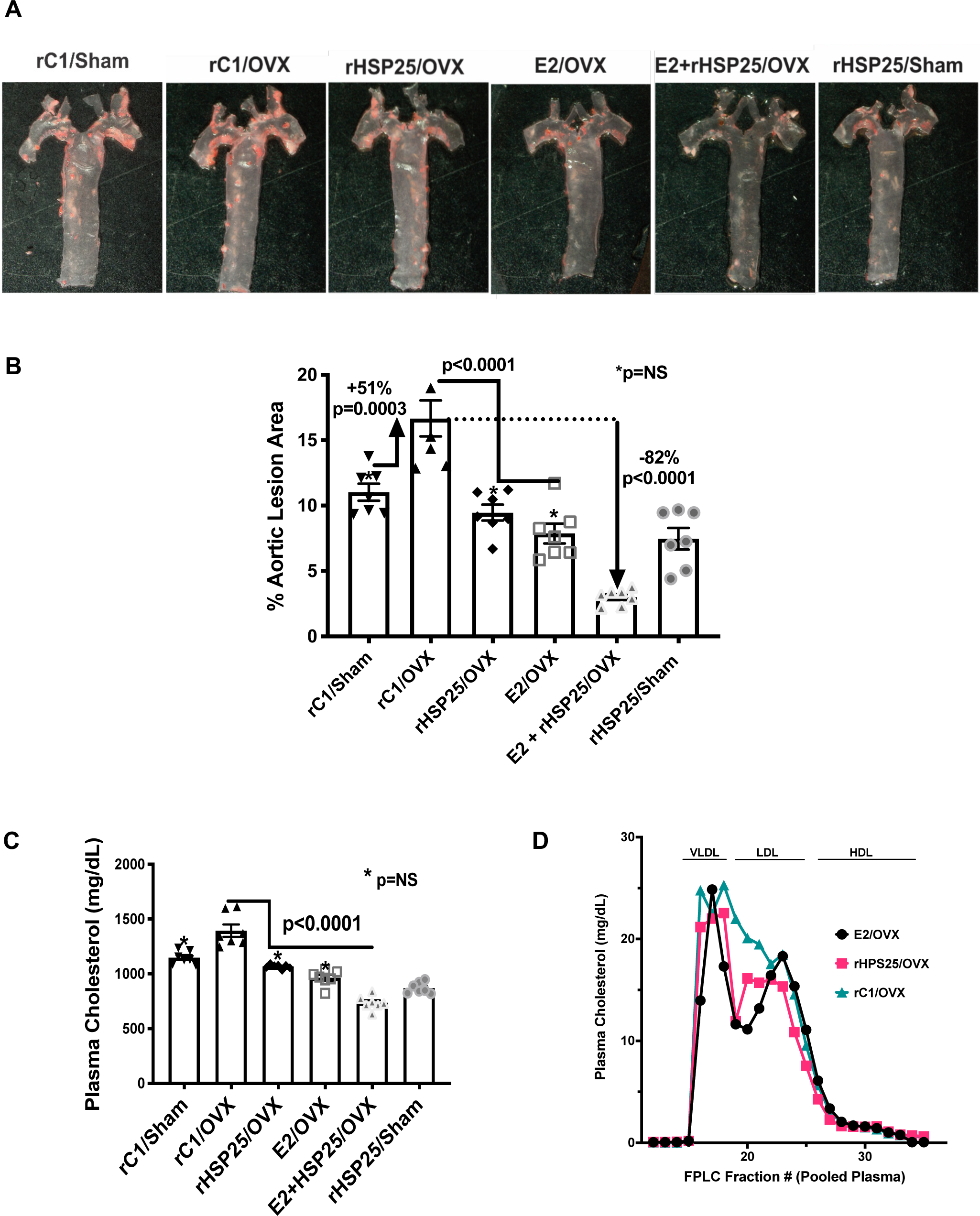
Atherosclerosis and Hyperlipidemia after surgical MP. A) Photomicrographs of *en face* thoracic aortae stained with Oil Red O. B) Quantitation of aortic atherosclerotic burden, as reflected by Oil Red O staining. Compared to the rC1 sham group there was an increase in aortic lesion area with rC1/OVX. Separately, rHSP25 and E2 reduced aortic atherosclerotic burden compared to rC1/OVX, but the largest decrease was noted with combined [E2+RHSP25] therapy. The atherosclerotic burden was lower in the rHSP25/sham *vs*. rC1/sham mice (p=0.04). C) OVX resulted in an increase in total plasma cholesterol in the rC1 mice. Compared to rC1/OVX, cholesterol levels were lower post-OVX with either rHSP25 or E2 therapies, and lowest with combined [rHSP25+E2] treatment. Cholesterol levels in the rHSP25/sham *vs*. rC1/sham mice (p<0.0001). D) Cholesterol measured in FPLC plasma fractions from pooled blood samples from rC1/OVX, rHSP25/OVX and E2/OVX group *ApoE*^*-/-*^ mice (n=8 per group). Lipid sub-fractions based on size are indicated above the cholesterol graphs. Compared to the rC1/OVX mice, there are reductions in the LDL subfraction in both rHSP25/OVX and E2/OVX groups. All statistical analyses used a one-way ANOVA with Tukey’s multiple comparison’s test.

### Murine Plasma Cholesterol Profiles and Lipoprotein Fractions

Total plasma cholesterol levels were determined using an enzymatic colorimetric assay kit (Wako Pure Chemical Industries, Ltd, 439-17501) (**Fig. 2C, Supplemental Fig. 1D, 2C, 2D**). The lipoprotein cholesterol distribution profile (VLDL, LDL, HDL) was evaluated in pooled plasma samples under fasting conditions after fractionation by fast protein liquid chromatography (FPLC) gel filtration on a single Superose 6 column (**Fig. 2D**).

### LDLR and PCSK9 Gene Expression in Murine Liver Tissue

As previously described, RNA was isolated from mouse liver tissue and used for Real-time PCR analysis of LDLR and PCSK9 gene expression (**Fig. 3A, 4B**).^16^ The specific murine primer sequence pairs, including those for the control gene, hypoxanthine-phosphoribosyltransferase (hHPRT1), are provided in **Supplemental Table 1**. RNA expression is presented as log fold change.

**Table 1.**

| Parameter | rHSP25 | E2 | E2 + rHSP25 |
| --- | --- | --- | --- |
| **Atherogenesis\*** | -43% | -52% | -82% |
| **Plasma Cholesterol\*** | -23% | -30% | -47% |
| **Anti-HSP25 Antibody** | Elevated | No change | Elevated |
| **LDLR Expression** | Upregulated mRNA (+219%)* & protein (+50%)* | No change | Upregulated mRNA (+165%)** & protein (51%)** |
| **PCSK9 Plasma\*** | -22% | +52% | +17% (-23%)** |
| **PCSK9 Hepatic mRNA\*** | No change | +148% | No change |
| **PCSK9 Promoter Activity\*\*\*** | -64% *** | +303% | +144% (-43%)** |
| • relative to rC1/OVX<br>** relative to E2 alone<br>*** [rHSP27 + PAb] treatment of HepG2 cells (PCSK9 promoter assay) |  |  |  |

**Fig. 3.**
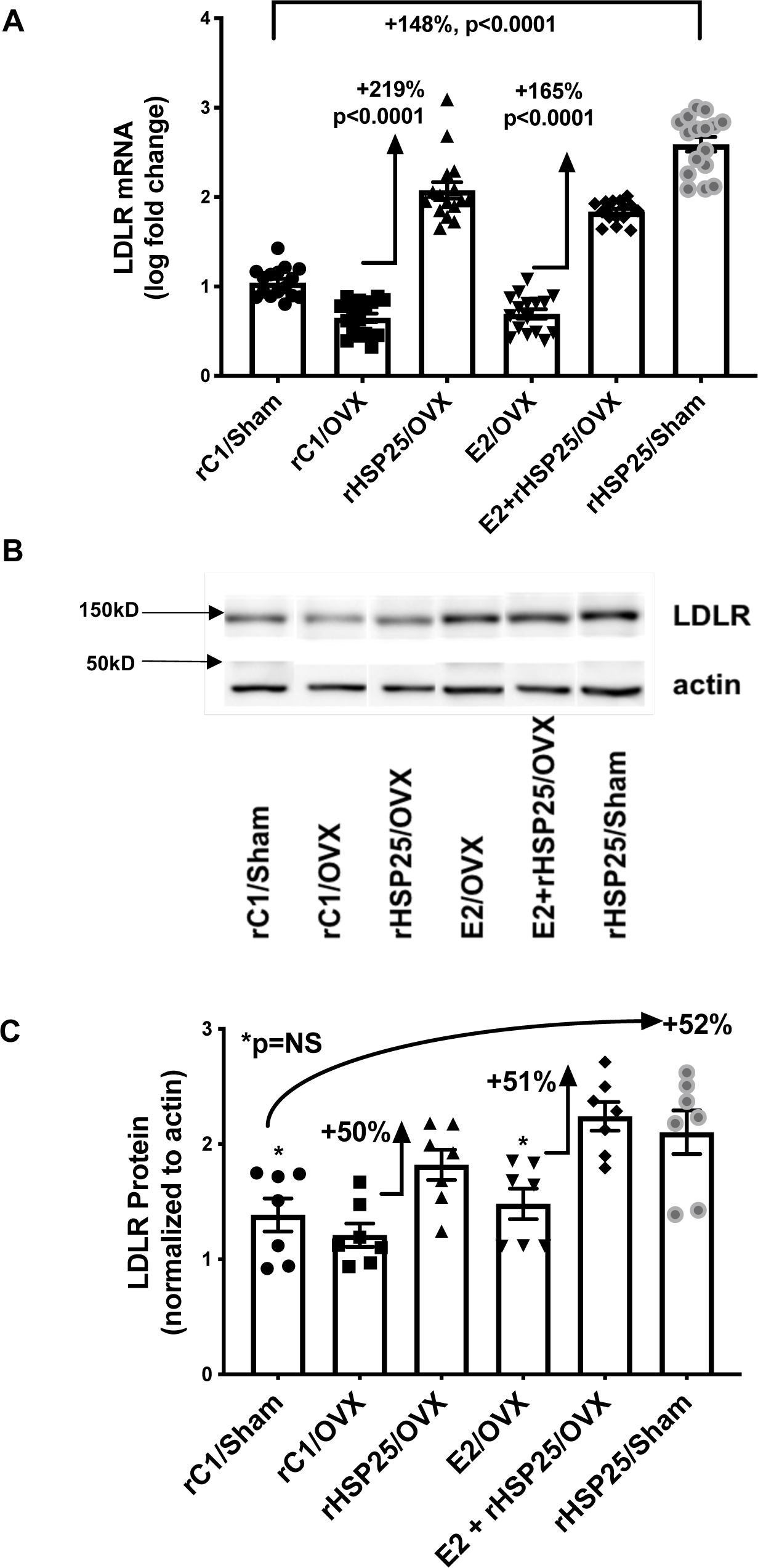
rHSP25 and E2 differentially regulate of LDLR. A) Compared to rC1/OVX mice, hepatic LDLR mRNA increased dramatically in the rHSP25/OVX but not the E2/OVX mice. In the OVX mice, adding rHSP25 to E2 noticeably increased hepatic LDLR mRNA expression compared to the E2/OVX group. There was a marked increase in LDLR expression in the rHSP25/sham *vs*. the rC1/sham mice. B) Hepatic LDLR protein expression in the OVX experiment involving *ApoE*^*-/-*^ mice, assessed by WB. Each LDLR protein band was normalized using β-actin as the loading control. C) LDLR protein expression was similar in the rC1/sham, rC1/OVX and E2/OVX mice, but appreciably increased with rHSP25 vaccination or combined [rHSP25 + E2] treatment post-OVX. LDLR protein expression was similarly increased in the rHSP25/sham *vs*. rC1/sham mice. All statistical analyses used a one-way ANOVA with Tukey’s multiple comparison’s test.

### LDLR Protein Expression in Murine Liver Tissue

Protein was harvested from the murine livers for SDS-PAGE. Multiple Western blots (WBs) were obtained for each protein assayed, and representative data are shown in **Fig. 3B, 3C**. Mouse liver tissue (200-300 mg) was placed inside a Precellys® homogenizing tube with 4 glass beads and 500 µL of RIPA lysis buffer (50 mM Tris HCL pH 7.4, 1% Triton X-100, 0.5% Sodium Deoxycholate, 150 mM NaCl, 2mM EDTA, protease inhibitor cocktail). Tissue was homogenized during two 30 s cycles at 6,000 RPM. The lysate was transferred to 2 mL tubes and further processed using a handheld sonicator at 50% amplitude for two 3 s pulses, centrifuged 45 mins at 16,000×g, and the supernatant layer below the fat and above the pellet was transferred to a new tube. Samples were centrifuged again for 10 mins at 16,000×g and the supernatants were transferred to new tubes. Protein lysates were quantified using a Bio-Rad DC Protein Assay, with equal amounts of protein (40-80 µg per well) loaded onto 10% Tris-Glycine gels for SDS-PAGE before being transferred to 0.2 μm nitrocellulose membranes ON at 4°C. Membranes were blocked in 5% Milk in TBS-Tween for 1h, washed, incubated in fresh primary antibodies with 5% bovine serum albumin (BSA) ON at 4°C, washed and incubated in secondary antibody before enhanced chemiluminescence detection using an ImageQuant LAS4000 imager (GE Healthcare). The primary antibodies used for WBs prepared using murine liver lysates were rabbit anti-LDLR (1:5,000; Abcam ab52818) and mouse anti-β-actin (1:10,000; sc-47778, Santa Cruz Biotechnology). The secondary antibodies used for WB were: anti-rabbit IgG HRP (Abcam ab97051) and anti-mouse IgG HRP (Jackson Immunoresearch 115-035-062).

### PCSK9 Measurement

Plasma PCSK9 levels (**Fig. 4A**) were measured using an ELISA kit specific to murine PCSK9 (R&D, DY3985) as per manufacturer’s instructions. Samples were diluted 1:100 in 1% BSA and a PCSK9 standard curve was generated for each experiment.

**Fig. 4.**
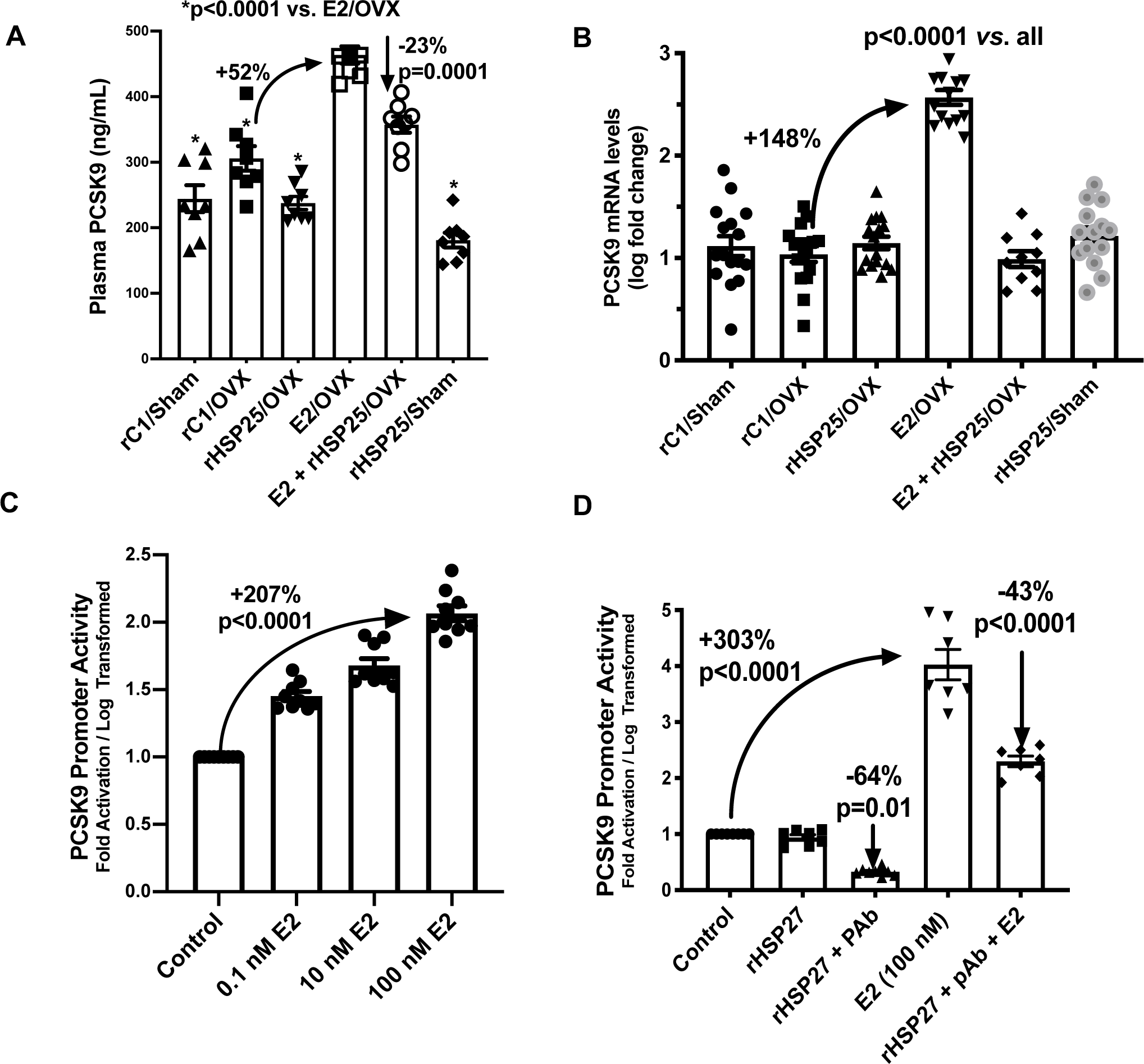
rHSP25 and E2 differentially regulate of PCSK9. A) Plasma PCSK9 protein levels increased with E2 treatment in the OVX mice, an effect that was attenuated when co-treated with rHSP25. rHS25 vaccination (alone) had no appreciable effects on plasma PCSK9 levels with or without OVX. B) Hepatic PCSK9 mRNA expression dramatically increased with E2 treatment (alone), but did not change when co-treated with rHSP25. Hepatic PCSK9 mRNA levels were unchanged in rHSP25 vaccinated mice and were unaltered by OVX. C) PCSK9 promoter activity in HepG2 cells was strongly upregulated by E2 in a dose dependent manner. D) Compared to control, rHSP27+PAb significantly reduced PCSK9 promoter activity, while E2 markedly upregulated PCSK9 promoter activity – an effect that was partially reduced by adding treatment with rHSP27. All statistical analyses used a one-way ANOVA with Tukey’s multiple comparison’s test.

**Fig. 5.**
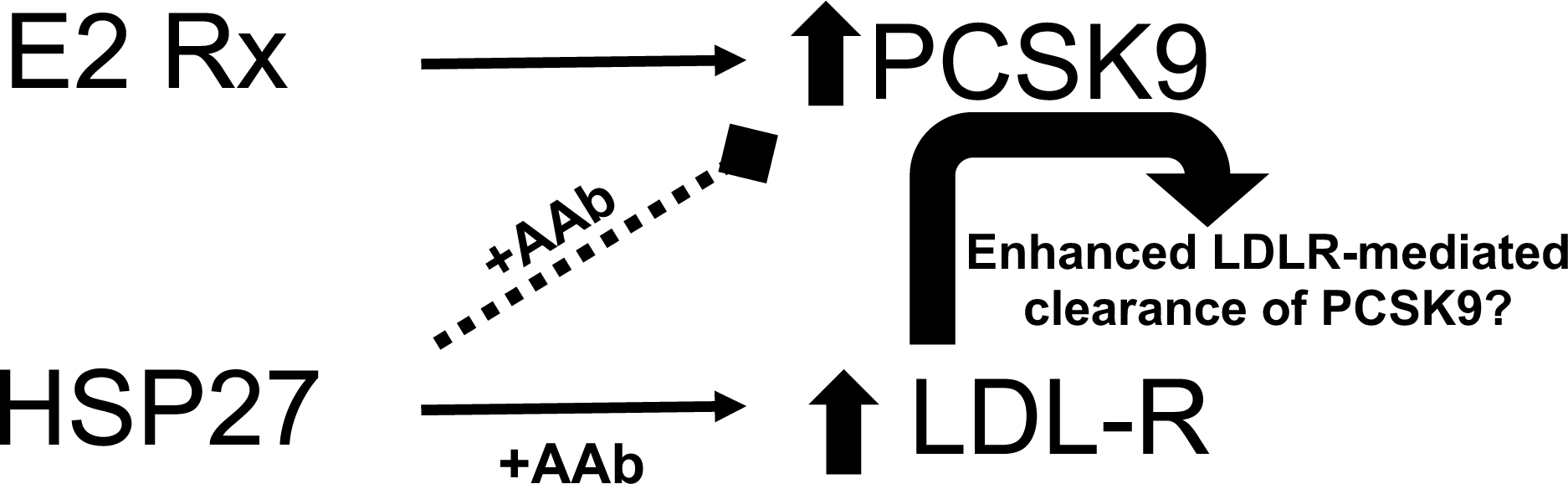
Graphical Abstract. Exogenous estrogen therapy increases PCSK9 expression. HSP27 vaccination upregulates LDLR expression and either directly or indirectly (via LDLR mediated clearance) reduces PCKS9 levels.

### PCSK9 Promoter Activity Assays

Cells from the human hepatic HepG2 cell line were grown in complete media (with 10% FBS and antibiotics) and seeded ON at 4×10^4^ cells/well in a 48-well culture plates, then: i) transiently transfected with a PCSK9 promoter firefly luciferase reporter construct, incorporated in the plasmids pGL3-PCSK9-D1 ^17^, and ii) co-transfected with a transfection control vector (pBLSV40 plasmid constitutively expressing a *Renilla* luciferase; generously donated by Dr. David Proud University of Calgary, Calgary, AB), so that the dual luciferase activities could be measured. The stoichiometry was: 500 ng of the PCSK9-D1 promoter construct and 25 ng of pBLSV40 vector, using 1 μL of Lipofectamine®2000 Transfection Reagent (ThermoFisher). Cells were then treated for 24 hrs with recombinant proteins (rHSP27, rC1) with or without a previously characterized and validated polyclonal anti-HSP27 IgG antibody (PAb) ^13^. PCSK9 promoter-driven firefly luciferase activity and control renilla luciferase activity were measured using the Firefly & Renilla Luciferase Single Tube Assay Kit (#30081-1; Biotium, Fremont, CA) according to the manufacturer’s protocol, with some minor modifications. Briefly, cells were washed with PBS then lysed in 110 μL of passive lysis buffer. Firefly luciferase activity was measured first by combining 30 μL of cell lysate with 50 μL of firefly working solution in a 12×75mm round-bottom tube, measuring luminescence then adding 5 μL of *Renilla* working solution to the same tube and again measuring luminescence. Luminescence was detected using a Monolight 3010 Luminometer (BD Biosciences). Absolute firefly luciferase activity was normalized against *Renilla* luciferase activity to correct for transfection efficiency. Triplicate wells were assayed for each transfection condition, and 3 independent transfection assays were performed for each reporter construct. The data was log transformed in order to facilitate parametric statistical analyses (**Fig. 4C, 4D**).

### Statistical analysis

For all data, continuous variables are expressed as mean ± standard error of the mean (SEM). Statistical significance was assessed by using a one-way ANOVA (Graph Pad Prism 8, La Jolla, CA, USA) and Tukey’s comparison of the groups. The direct comparison of unpaired treatment groups was performed using a Student’s t-test. Quantitative PCR gene expression data were log-transformed and are reported as log-fold change in order to more normally distribute the data and permit parametric statistical analyses. A p-value < 0.05 was considered significant.

## Results

### a) Overview of rHSP25 Vaccination *vs*. E2 Therapy Experiment in OVX mice

To test the hypothesis that rHSP25 vaccination reduces atherogenesis in OVX mice, we subjected atherosclerosis-prone *ApoE*^*-/-*^ mice to OVX, followed by 6 weekly subcutaneous injections with rHSP25 with the adjuvant Alum (**Fig. 1A, 1B**). Another group of mice underwent a sham surgery (the ovaries were not removed) followed by rHSP25 vaccination. For comparison with the clinical scenario of hormonal therapy for MP, a group of OVX *ApoE*^*-/-*^ mice underwent subcutaneous implantation of a slow release E2 pellet, while another group received both E2 and rHSP25 vaccination. The inactive truncated C-terminus of HSP27, rC1 (mixed with Alum), was administered as a negative control to OVX or sham surgery mice. All mice (7-8 per group) received a HFD for 5 weeks, beginning one week after the initiation of therapies.

### b) Validation of the Vaccination response

The rise in IgG anti-HSP25 levels were similar for all mice vaccinated with rHSP25 whether it was combined with E2 or not (**Fig. 1C**). The antibody response to vaccination with rC1 was similar to that with rHSP25, thereby indicating that rC1 remained immunogenic. E2 or PBS treated mice had very low (background) levels of AAbs.

### c) Validation of the Murine Model of Menopause and Physiological Estrogen Therapy

We confirmed that this OVX model was appropriate for the study of MP atherogenesis by showing that OVX increased atherosclerotic burden and cholesterol levels. *ApoE*^*-/-*^ mice subjected to OVX had more aortic atherosclerosis than sham operated mice [e.g., PBS sham *vs*. PBS OVX: 45% increase (11.8 ± 1.2 vs. 17.1 ± 1.7; respectively, p=0.03); and rC1 OVX compared to rC1 sham: 65% increase (18.2 ± 1.9 *vs*. 11.0 ± 0.7; respectively, p=0.006)] **(Supplemental Fig. 1B, 1C)**. Similarly, total plasma cholesterol levels 1-day pre-euthanasia were higher in OVX mice compared to sham operated mice [e.g., PBS/sham compared to PBS/OVX: 23% increase (1,119 ± 32 *vs*. 1,374 ± 34; respectively, p=0.0001), and rC1/sham compared to rC1/OVX: 21% increase (1,136 ± 23 *vs*. 1,369 ± 55; respectively, p=0.002)] **(Supplemental Fig. 1D)**. As measurement of estrogen levels in mice is technically challenging, we confirmed that the estrogenic therapy in the E2 pellet (0.1 mg/60 day) provided a physiological level of hormone replacement by examining the uteri at the end of the experiment. The uterine weights of the E2 treated mice were similar to those of the sham mice, thereby confirming that the E2 dose was likely in a physiological range (**Fig. 1D, 1E**). In contrast, the rHSP25/OVX uteri were similar to those of the rC1/OVX mice showing 4.5-fold atrophy *vs*. rC1/sham mice uteri; suggesting that rHSP25 had no direct or indirect estrogenic properties.

### d) Validation of rC1 as a Negative Control Treatment in OVX / Atherogenesis Model

Four control groups (PBS/sham, rC1/sham, PBS/OVX and rC1/OVX) were used to assess the validity of using rC1 as a negative control. rC1 is an excellent control treatment as it is prepared in an identical manner as rHSP25 using recombinant technology in *E. coli*. The degree of atherosclerosis was similar for the sham operated mice that received PBS *vs*. rC1 (11.8 ± 1.19 *vs*. 11.03 ± 0.65 respectively; p=0.58), as well as the OVX mice that received PBS *vs*. rC1 (17.1 ± 1.67 *vs*. 18.2 ± 1.93; respectively, p=0.68) (**Supplemental Fig. 2A, 2B**). Likewise, cholesterol levels were similar amongst the PBS sham *vs*. rC1 sham mice (1,119 ± 31.64 *vs*. 1,136 ± 22.83; respectively, p=0.67) and the PBS OVX *vs*. rC1 OVX mice (1,374 ± 33.75 *vs*. 1,369 ± 54.88; respectively, p= 0.94) (**Supplemental Fig. 2C, 2D**). Therefore, for the purpose of comparing the effect of the active treatment groups (rHSP25 or E2), the rC1/OVX group was used as the reference group for the effect of OVX, as the degree of atherogenesis as well as the cholesterol levels were similar to those found in the PBS/OVX cohort.

### e) rHSP25 Vaccination and E2 Therapy Post-OVX are Comparable for Attenuation of Atherogenesis and Lowering Cholesterol

Mice subjected to OVX alone (rC1/OVX) had 51% more aortic atherosclerosis compared to non-OVX (rC1/sham) mice (16.7 ± 1.4 *vs*. 11.0 ± 0.6; **Fig. 2A, 2B)**. The atherosclerotic burden was similar in the rHSP25/OVX and E2/OVX mice, and markedly reduced compared to rC1/OVX (−43% and -52%, respectively; p<0.0001). Interestingly, the treatment of OVX mice with combination of [rHSP25 + E2] showed the lowest atherosclerotic burden (−82% *vs*. rC1/OVX; p<0.0001). Aortic lesion formation was attenuated in the rHSP25/sham *vs*. rC1/sham mice (p=0.04).

While the baseline cholesterol levels in all mice fed a normal chow diet were similar (515 ± 82 mg/dL), by the second week of the HFD the levels peaked and then fell by varying degrees over the ensuing 3 weeks of the HFD. Terminal plasma cholesterol levels (**Fig. 2C**) were measured 1-day prior to euthanasia, with the highest levels found in the rC1/OVX group (1,393 ± 57 mg/dl), which was 21% higher (p<0.0001) than the rC1/sham mice (1,148 ± 23 mg/dl). Both the rHSP25/OVX and E2/OVX mice had similar cholesterol levels that were lower than the rC1/OVX mice (1,068 ± 8 or -23%, and 964 ± 27 or -30%, respectively; both p<0.0001), while the lowest levels were found in the combined therapy [rHSP25 + E2/OVX] mice (736 ± 24 or - 47%; p<0.0001 *vs*. all OVX groups). Cholesterol levels were lower in the rHSP25/sham compared to rC1/sham mice (p<0.0001). As demonstrated by FPLC, the LDL-C fraction was reduced in both the E2/OVX and rHSP25/OVX groups relative to the rC1/OVX mice (**Fig. 2D**).

### f) Unlike rHSP25 Vaccination, E2 Therapy Did not Increase LDLR Expression Post-OVX

rHSP25 alone or in combination with E2 markedly increased hepatic LDLR mRNA expression (+219% for the rHSP25/OVX *vs*. rC1/OVX and +165% for the [rHSP25 + E2] *vs*. E2/OVX mice; p<0.0001 for both; **Fig. 3A**). LDLR protein expression also increased in the mice vaccinated with rHSP25 (50% for the rHSP25/OVX *vs*. rC1/OVX, 51% for the [E2 + rHSP25/OVX] *vs*. E2/OVX; p=0.04 and p=0.006, respectively; **Fig. 3B, 3C**). Surprisingly, E2 therapy in isolation had no effect on LDLR mRNA or protein expression.

### g) E2 but Not rHSP25 Vaccination Increases PCSK9 Gene Expression

Plasma PCSK9 levels were much higher in E2/OVX (464 ± 13 ng/ml) compared to the other four treatment groups (*p*<0.0001) – including a 52% increase relative to rC1/OVX (**Fig. 4A**). Combining [rHSP25 + E2] therapy post-OVX resulted in a 23% reduction in PCSK9 levels (357 ± 12 ng/ml; p=0.0001) relative to E2/OVX alone. Plasma PCSK9 levels were 26% lower in the rHSP25/sham *vs*. rC1/sham mice (p=0.046). Hepatic PCSK9 mRNA expression was 148% higher in the E2/OVX compared to rC1/OVX and all other murine treatment groups (p<0.0001) (**Fig. 4B**).

The PCSK9 promoter is thought to contain two estrogen response elements (EREs), as predicted *in silico* using the MatInspector analysis tool (optimized matrix threshold = 0.9 for both; https://www.genomatix.de/online_help/help_matinspector/matinspector_help.html). Hence, we assessed PCSK9 promoter activity in HepG2 cells treated for 24 hrs with escalating concentrations of E2 (0.1, 10, 100nM) and noted a robust dose-dependent increase in activity (+207% log transformed increase with 100 nM E2, p<0.0001) (**Fig. 4C**). Likewise, treating HepG2 cells for 24 hrs with rHSP27 alone (100 μg) had no effect, yet there was a 64% decrease in PCSK9 promoter activity (p=0.01) when cells were treated with [rHSP27 + PAb] (**Fig. 4D**). Moreover, there was a remarkable 303% fold increase (log transformed) in PCSK9 promoter activity with E2 treatment (100 nM; p<0.0001) that fell by 43% (p<0.0001) with [rHSP27 + PAb] treatment.

## Discussion

Using a surgical model of MP in atherosclerosis-prone ApoE^-/-^ mice fed a HFD we demonstrate that rHSP25 vaccination, like E2 therapy, is atheroprotective. As well, we confirmed a recent observation ^13^ that rHSP25 vaccination promotes the upregulated expression of the LDLR – an effect that is not associated with E2 therapy. Unlike rHSP25 vaccination, E2 therapy is associated with increased PCSK9 synthesis and plasma levels (**Fig. 4A, 4B**), and showed a dose dependent increase in PCSK9 promoter activity that was partially abrogated with concomitant treatment with [rHSP27 + PAb] (**Fig. 4C, 4D**).

The specific mechanisms that account for the relative protection against atherosclerotic complications in pre-MP women are poorly understood, yet there is a common belief that one of the main cardiovascular benefits of estrogens is the lowering of LDL-C due to an increase in LDLR expression. However, the pre-clinical and clinical literature suggest that the effects of estrogens on LDLR vary. For example, in rats LDLR activity and expression are reduced with OVX but improve with estrogen therapy ^18, 19^. But, in non-OVX mice treated with (even) supra-physiological E2 concentrations, LDLR expression does not change ^20, 21^. As the *ApoE*^*-/-*^ mice in our study did not show an increase in LDLR with E2, this does not seem to be due to an inherent problem with LDLR expression – as these same mice had a robust increase in LDLR expression in response to rHSP25 vaccination. Moreover, the fact that E2 treatment is atheroprotective in *LDLR*^*-/-*^ mice also points to estrogenic effects that may be at a distance from LDLR modulation 22. The evidence that E2 modulates LDLR expression in human hepatocytes is also far from convincing ^23, 24^. While there is indirect evidence of increased LDLR binding activity in liver biopsy tissue from two male patients receiving estrogen therapy for prostate cancer the estrogenic compounds administered to these men were not in a physiological range, and did not lower their cholesterol levels ^25^. Given that >1,000 human hepatic genes display a sex-bias in their expression, particularly those known to participate in or regulate lipid metabolism, and that the LDLR gene is more highly expressed in the female liver, experimental results in male subjects should be interpreted with caution ^26^.

PCSK9 was discovered less than 2 decades ago, and there is a lot to learn about its transcriptional regulation, including the potential role of two estrogen response elements in its promoter. The estrogen mediated upregulation of PCSK9 expression noted in the current studies fits with clinical observations showing that: i) PCSK9 levels are higher in pre-MP women than men ^27^, and ii) PCSK9 levels are markedly elevated in pregnant compared to non-pregnant age-matched women (493 *vs*. 290 ng/ml) ^28^. Indeed, in the third trimester of pregnancy E2 levels increase 25-50 fold and total cholesterol levels rise 50-75% ^29^. Whether the surges in E2 levels that occur during the normal menstrual cycle or with exogenous estrogenic therapies also modulate PCSK9 levels remains to be studied. Certainly, we have more to learn about PCKS9 transcriptional regulation and estrogenic hormones, including the potential differences between estrogen receptor alpha *vs*. beta modulators. Moreover, are there yet to be discovered important physiological perturbations in women who are treated with therapies targeting PCSK9? Perhaps estrogens modulate PCSK9 levels for reasons other than cholesterol metabolism, as we are only now beginning to understand the diverse biological roles of PCSK9 ^30^. Given the relative paucity of sex-disaggregated analyses of PCSK9 inhibitor clinical trials, this will be an important question to address in the future.

One study that does not fit with our observations on the role of E2 in upregulating PCSK9 levels is the Dallas Heart Study, which found that PCSK9 levels are higher in post-*vs*. pre-MP women, yet no difference in PCSK9 levels between women receiving estrogenic compounds or not ^27^. Unfortunately, it is not known if the higher PCSK9 levels in the older post-MP women are clearly related to loss of ovarian function or if the estrogenic therapies that they received were physiological. Indeed, at least three other studies suggest that PCSK9 levels simply correlate with age ^31-33^.

In summary, although we did not see major differences in atheroprotection with E2 *vs*. rHSP25 therapy *in vivo*, we uncovered some subtle mechanistic differences that require further consideration – particularly the E2-induced increase in PCSK9 expression. We summarize the results of our studies in the **Table 1** and the **Graphical Abstract**. Briefly, E2 upregulates PCSK9 expression and plasma levels, whilst this is not true for rHSP25 vaccination. As we recently reported, rHSP25 vaccination is associated with a strong increase in LDLR expression.

Hence, our working hypothesis is that the decrease in PCSK9 levels in the rHSP25 vaccinated mice is (at least in part) due to increased clearance of PCSK9 by (the more abundant) LDLRs shuttling it for intracellular disposal. However, this does not appear to be the sole mechanism, as rHSP25 vaccination also produced a reduction in PCSK9 promoter activity – an observation that will require more refined study at the promoter level. Currently, we are trying to better comprehend how HSP27 (or HSP25) alone or in combination with its autoantibody, modulate LDLR and PCSK9 transcription. We do know that in the presence of AAbs, HSP27 shows enhanced docking at the cell membrane, interacts with key receptors and signaling pathways, and can undergo internalization ^12^. As low HSP27 and AAb levels are associated with atherosclerosis,^13^ we want to develop immunotherapies that boost AAb levels in order to potentiate the biological activity of existing (low) levels of HSP27, especially in post-MP women.

## Abbreviations

AAb: anti-HSP27 (or anti-HSP25) antibodies
ApoE^-/-^: apolipoprotein E null
a.u.: absorbance units
E2: 17β estradiol
HFD: high fat diet
HPRT: hypoxanthine-phosphoribosyltransferase
HSP27: Heat Shock Protein 27
HSP27^o/e^: HSP27 over-expressing
LDL-C: low density lipoprotein cholesterol
LDLR: low density lipoprotein receptor
MP: menopause
mRNA: messenger RNA
NF-κB: nuclear factor kappa light chain enhancer of activated B cells
ON: overnight
OVX: ovariectomy
PAb: polyclonal anti-HSP27 IgG antibody
PBS: phosphate buffered saline
PCSK9: Proprotein Convertase Subtilisin/Kexin type 9
rC1: recombinant C-terminal truncation of HSP27 spanning amino acids 90-205
rHSP27: recombinant HSP27
SDS-PAGE: sodium dodecyl sulfate and polyacrylamide gel
WB: Western Blot

## Acknowledgments

We are indebted to the Libin Cardiovascular Institute of Alberta and its community partners for their support, as well as the staff in the University of Calgary Cumming School of Medicine Health Sciences Animal Resource Centre.

## Author Contributions

N. Maarouf and E.R. O’Brien designed the studies. N. Maarouf performed all *in vivo* experiments. N. Maarouf, Y.-X. Chen, C. Shi, J. Deng, C. Diao, M. Rosin, V. Shrivastava performed or contributed to some of the *in vitro* and *ex vivo* research; N. Maarouf, C. Diao and J. Liu performed or contributed to the design and analysis of the promoter studies; N. Maarouf, V. Shrivastava, Z. Batulan and E.R. O’Brien analyzed all other data; N. Maarouf and E.R. O’Brien wrote the manuscript.

