## Supplemental Figures for "Heat Shock Protein 27 versus Estrogen Therapy for Post-Menopausal Atherosclerosis: Rethinking Mechanisms of Cholesterol Lowering"

### Slide 1
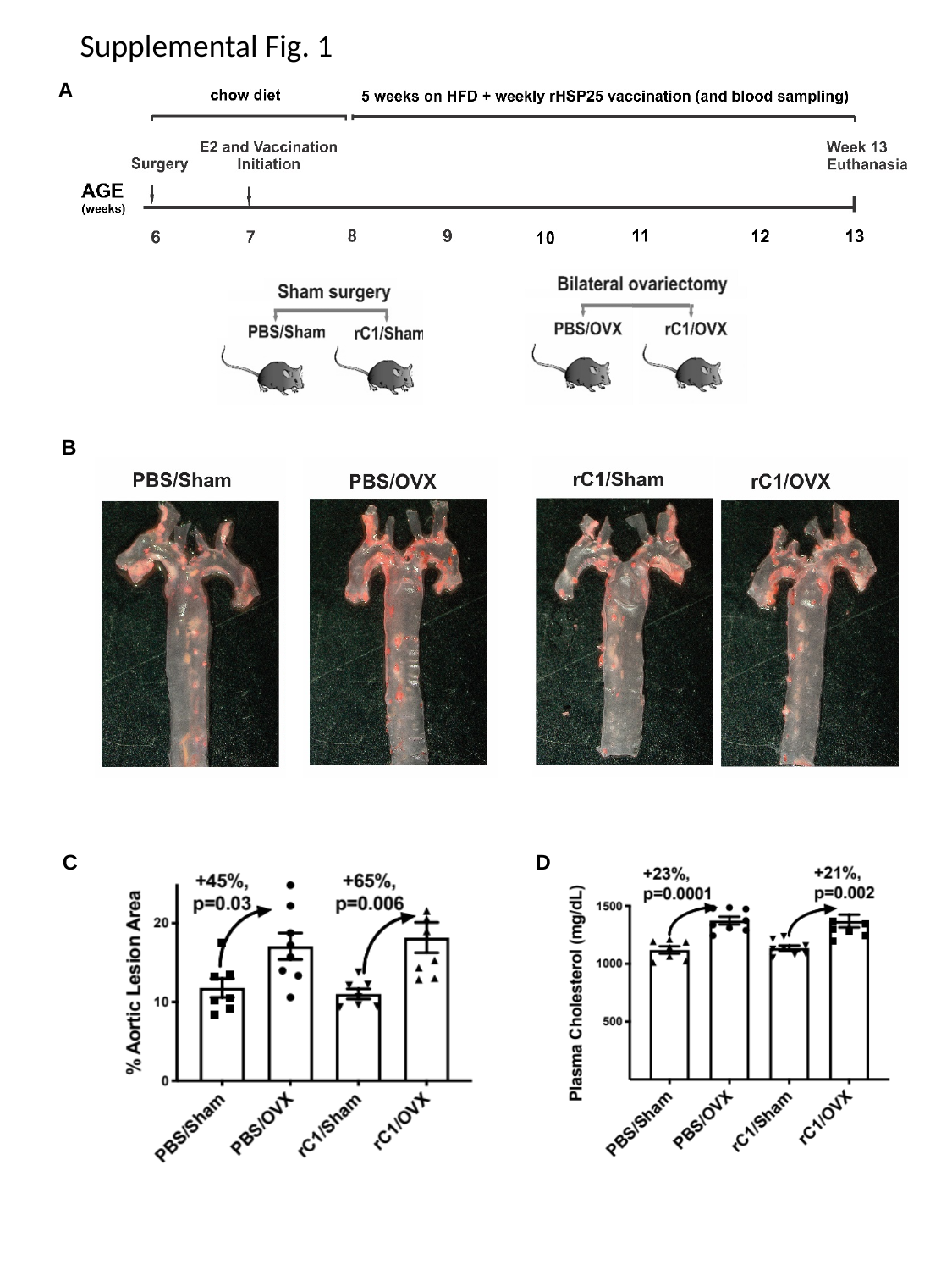

Supplemental Fig. 1
A
B
C
D

### Slide 2
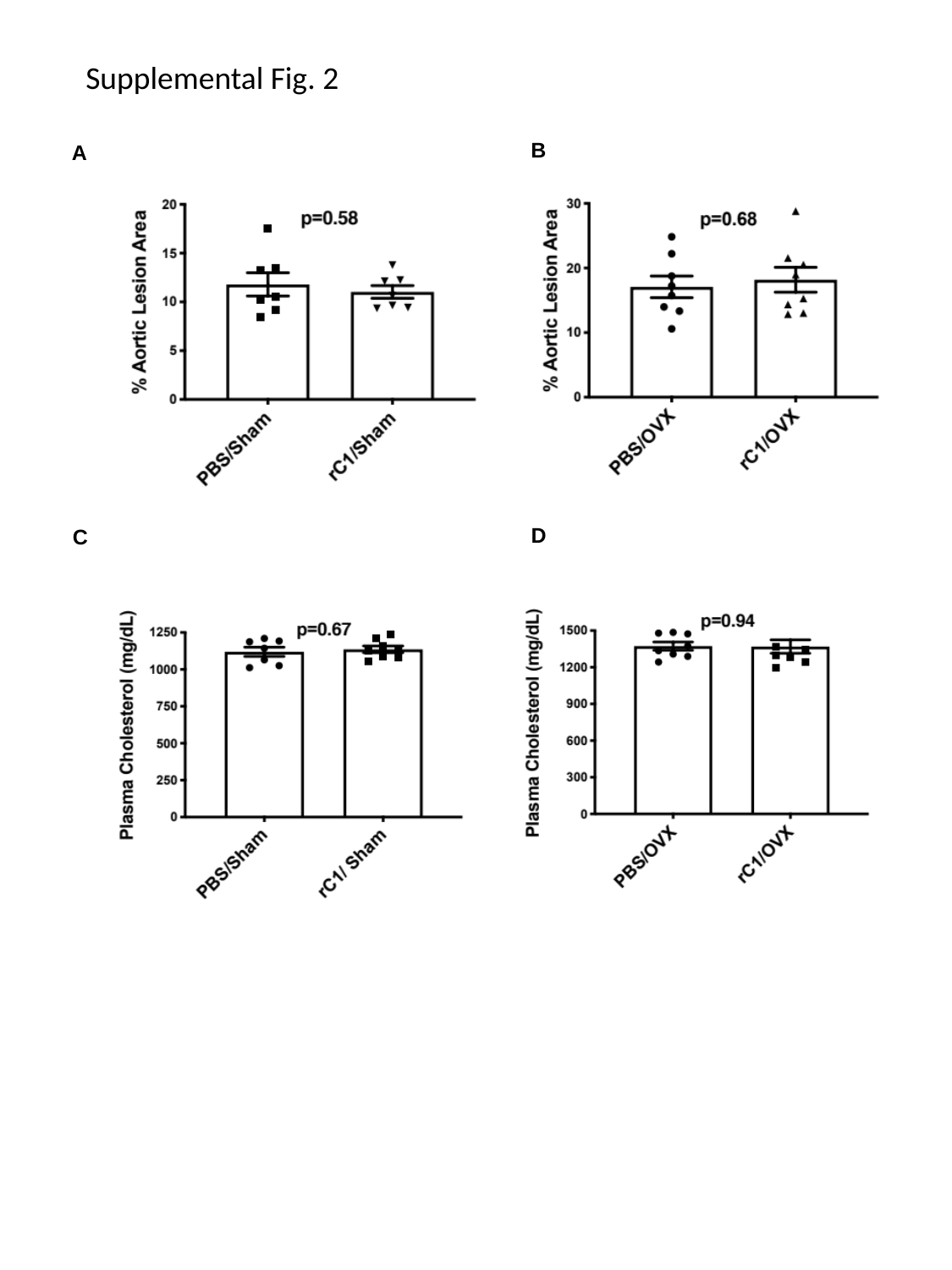

Supplemental Fig. 2
B
A
D
C

### Slide 3
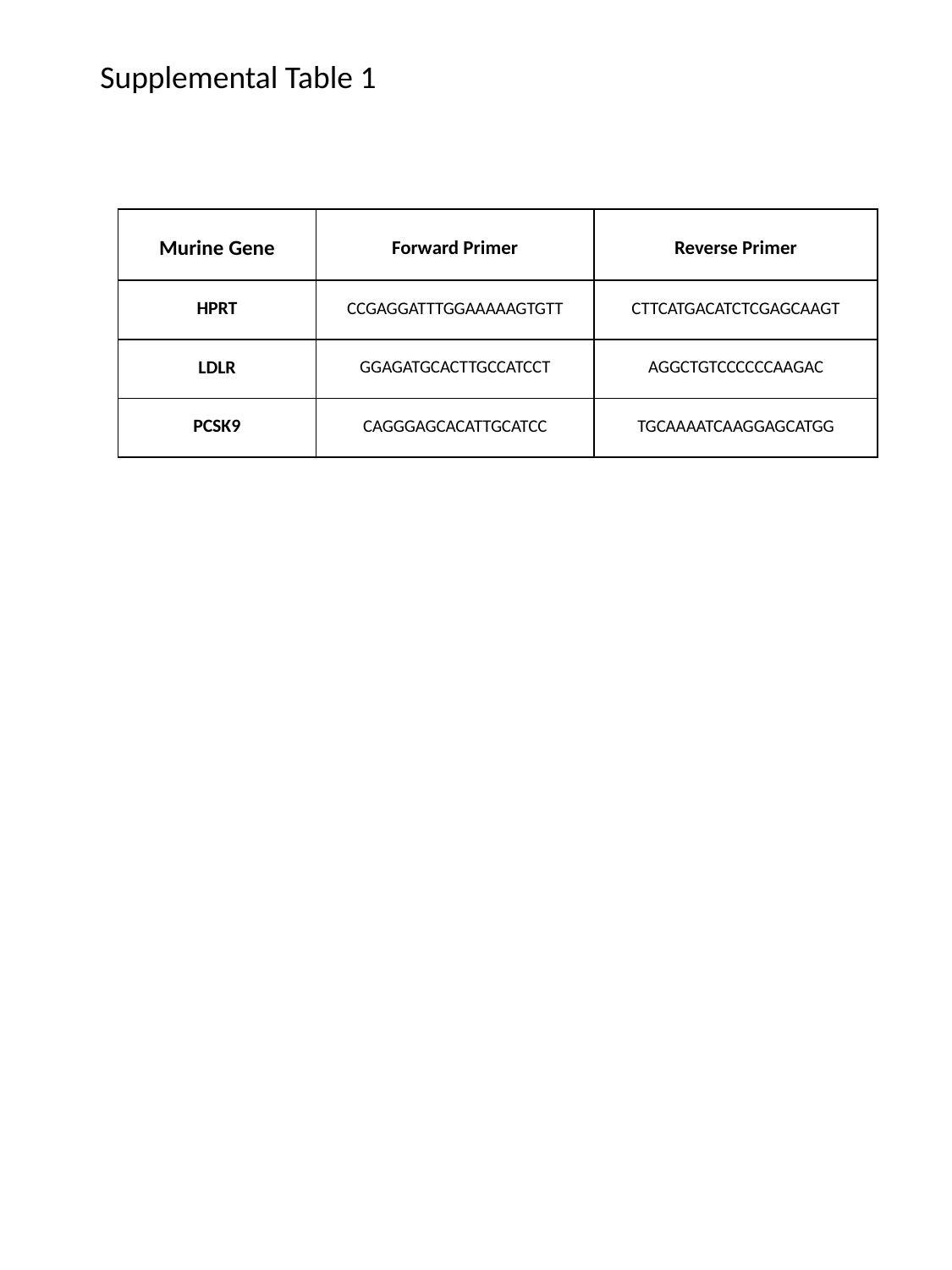

Supplemental Table 1
| Murine Gene | Forward Primer | Reverse Primer |
| --- | --- | --- |
| HPRT | CCGAGGATTTGGAAAAAGTGTT | CTTCATGACATCTCGAGCAAGT |
| LDLR | GGAGATGCACTTGCCATCCT | AGGCTGTCCCCCCAAGAC |
| PCSK9 | CAGGGAGCACATTGCATCC | TGCAAAATCAAGGAGCATGG |
