## Supplemental Figure Legends for "Heat Shock Protein 27 versus Estrogen Therapy for Post-Menopausal Atherosclerosis: Rethinking Mechanisms of Cholesterol Lowering"

**Supplemental Figures & Legend**

**Supplemental Fig. 1.** Control groups for the murine model of atherogenesis after OVX

A) Study outline: Age-matched, sexually mature female *ApoE^-/-^* mice were subjected to either OVX or a sham operation at 6 weeks of age. After one week of recuperation from the surgery, mice were randomized to begin the weekly active treatments (described in Fig. 1) or the control treatments, PBS or rC1 (shown here). One week later, these mice were switched to a HFD for 5 weeks until euthanasia.
B) Photomicrographs of aortic lesion burden in the *ApoE^-/-^* mice subjected to sham or OVX procedures and treated with PBS or rC1.
C) Quantitation of aortic lesion burden: In the subgroups of PBS- and rC1-treated mice OVX resulted in marked increases in the percentage aortic wall lesion area compared to sham mice.
D) Plasma cholesterol levels: In the subgroups of PBS- and rC1-treated mice, OVX resulted in moderate increases in plasma cholesterol levels compared to sham mice.

All statistical analyses used a one-way ANOVA with Tukey’s multiple comparison’s test.

**Supplemental Fig. 2.** Direct comparison of atherogenesis and cholesterol levels in control mice

A) Atherosclerotic lesion burden was similar in the PBS- *vs.* rC1-treated murine control groups that underwent a sham operation.
B) Atherosclerotic lesion burden was similar in the PBS- *vs.* rC1-treated murine control groups that underwent OVX.
C) Terminal total plasma cholesterol levels were similar in the PBS- *vs.* rC1-treated murine control groups that underwent a sham operation.
D) Terminal total plasma cholesterol levels were similar in the PBS- *vs.* rC1-treated murine control groups that underwent OVX.

All statistical analyses used an unpaired t-test.

**Supplemental Table 1.** Sequences of the qPCR primers used to assess expression of indicated murine genes.
